# Explainable AI reveals changes in skin microbiome composition linked to phenotypic differences

**DOI:** 10.1101/2020.07.02.184713

**Authors:** Anna Paola Carrieri, Niina Haiminen, Sean Maudsley-Barton, Laura-Jayne Gardiner, Barry Murphy, Andrew Mayes, Sarah Paterson, Sally Grimshaw, Martyn Winn, Cameron Shand, Will Rowe, Stacy Hawkins, Ashley MacGuire-Flanagan, Jane Tazzioli, John Kenny, Laxmi Parida, Michael Hoptroff, Edward O. Pyzer-Knapp

## Abstract

Alterations in the human microbiome have been observed in a variety of conditions such has asthma, gingivitis, dermatitis and cancer, and much remains to be learned about the links between the microbiome and human health. The fusion of artificial intelligence with rich microbiome datasets can offer an improved understanding of the microbiome’s role in our health. To gain actionable insights it is essential to consider both the predictive power and the transparency of the models by providing explanations for the predictions.

We combine the effort of collecting a corpus of leg skin microbiome samples of two healthy cohorts of women with the development of an *explainable artificial intelligence (EAI)* approach that provides accurate predictions of phenotypes and explanations. The explanations are expressed in terms of variations in the abundance of key microbes that drive the predictions.

We predict skin hydration, subject’s age, pre/post-menopausal status and smoking status from the leg skin microbiome. The key changes in microbial composition linked to skin hydration can accelerate the development of personalised treatments for healthy skin, while those associated with age may offer insights into the skin aging process. The leg microbiome signatures associated with smoking and menopausal status are consistent with previous findings from oral/respiratory tract microbiomes and vaginal microbiomes respectively. This suggests that easily accessible microbiome samples could be used to investigate health-related phenotypes, offering potential for non-invasive diagnosis and condition monitoring.

Our EAI approach sets the stage for new work focused on understanding the complex relationships between microbial communities and phenotypes. Our approach can be applied to predict any conditions from microbiome samples and has the potential to accelerate the development of microbiome-based personalised therapeutics and non-invasive diagnostics.

## Introduction

The associations between the human microbiome and individual health-related phenotypes are increasingly studied, with applications such as use of intervention strategies including prebiotics and probiotics(1), and personalized therapies(2). Improved understanding of how microbial taxa contribute to health and wellbeing, coupled with the increasing ability to measure the microbiome, can drive and accelerate the development of personalised microbiome-based treatments. As microbiome research expands and as sequencing technologies continue to advance, the volume and complexity of the data collected inexorably increases. Therefore, the necessity to develop sophisticated methods for analysing microbiome data to derive actionable insights becomes increasingly important. Machine learning (ML) has the potential for building predictive models that can provide powerful comprehensions into the complex interaction between microbial communities and their host organisms(3). The application of ML to microbiome data has answered several important questions regarding the clustering of microbial species, taxonomic assignment, comparative metagenomics, gene prediction.

Recently, a number of studies have been published on phenotype prediction from microbiome data(4–10), where some particularly focus on identifying discriminatory microbial taxa(11) or microbial signatures(12). As ML is becoming increasingly deployed in patient-relevant settings, it is essential to consider both the predictive power of the models and the transparency of the recommendations by providing an explanation for the predictions(13). This is often referred to as the *explainability* of the models. Recent work in this direction has demonstrated that interpretable AI/ML for microbiome data is increasingly applied as explanations for the predictions are needed (14)(15)(16). However, more can be done in terms of explaining the mechanisms of the ML algorithm predictions for microbiome data.

We propose an *explainable artificial intelligence* (EAI) approach to identify key predictive taxa and to investigate how the distributions of these microbial taxa drive the prediction of different phenotype values. The explanations are expressed in terms of key variations in microbiome composition. Our streamlined EAI approach includes three fine-tuned machine learning models – random forest (RF) (17), XGBoost (18) and neural network (NN) (19) – to predict the phenotype. We refer to the three models as a *bag of models*. Our approach explains the predictions using an explainable AI algorithm called SHapley Additive exPlanations (SHAP) (20). Obtaining explanations for samples in which the model is unreliable could lead to poor quality explanations. In order to achieve high-quality explanations, we repeatedly cross-validated predictions and explanations. Additionally, we propose a new approach of providing explanations for a sub-set of samples for which the ML model reliably predicted phenotypes. We refer to these samples as *exemplars*. Finally, our EAI approach creates consensus explanations using the exemplars derived from the three ML models. This process can be thought of as using the bag of models as a representation from which explanations are generated, while encouraging general explanations over those that are specific to one model. A full discussion of our EAI approach can be found in Methods.

To exemplify the power of our EAI approach as a means to derive actionable insights from complex interactions between microbes and their hosts, this study is focused on the human skin microbiome. The technical ease of acquiring skin microbiome data sets – it is reliant only upon swabbing or scrubbing of skin to collect samples – and the recent increase in interest in analysing the human skin microbiome(21–25) makes skin an appealing target. Human skin is in fact a large, heterogeneous organ that protects the body from pathogens while sustaining microorganisms that influence human health and disease(26).

For this study, a total of 1234 time-series leg skin microbiome samples (bacterial 16S rRNA gene sequencing) as well as associated skin hydration measures (visual assessment, pH, conductance, capacitance) were collected from 63 Canadian women (21-65 years of age) displaying healthy (non-dry) or moderately dry skin. The time series samples were collected between April and July 2017. Supplementary Figure 1 shows the number of samples per subject for the Canada cohort. Additional phenotype data (i.e., age, smoking and menopausal status) were obtained from subject questionnaires. This study is mainly focused on the Canada cohort. However, to validate and investigate the generalizability of the EAI approach we used a second independent UK cohort of 278 samples from 102 women as a true hold-out test set to predict skin hydration and age with the bag of model trained on the Canada cohort. We did not use the UK study for menopausal and smoking status prediction as the respective metadata were not available for this cohort. Table 1 and Supplementary Figure 1 report the distributions of the four phenotypes for the Canada and UK cohorts. See section Methods and Clinical design in Supplementary Information for more detail. The microbial features used by our EAI approach are derived from the observed genera, i.e., Operational Taxonomy Units (OTUs) at genus level. Therefore, we use the terms *genera* and *features* interchangeably. Relative abundances of 186 genera were obtained from taxonomic analysis of the sequenced reads of both studies.

**Table 1.** Metadata for Canada and UK cohorts. See Supplementary Figure 1 for more details on the distributions of corneometer, age and samples per subjects. Also see Supplementary Notes and Methods for more detail on the sample collection.

| Time series Samples | Country | Sex | Corneometer | Age | Menopausal status | Smoking status |
| --- | --- | --- | --- | --- | --- | --- |
| 1234 samples | Canada | Female | [2.4-57.2]<br>Median ~ 24.73<br>Mean ~24.67<br>Std ~8.87 | [21-65] Median ~56.0<br>Mean ~52.7<br>Std ~11.36 | 324 pre-menopausal<br>876 post-menopausal<br>34 peri-menopausal | 286 smoking<br>948 non-smoking |
| 278 samples | UK | Female | [8.1-78.4]<br>Median ~27.3<br>Mean ~31.70<br>Std ~12.42 | [19-55]<br>Median ~39.5<br>Mean ~38.74<br>Std ~9.09 | - | - |

**Figure 1.**
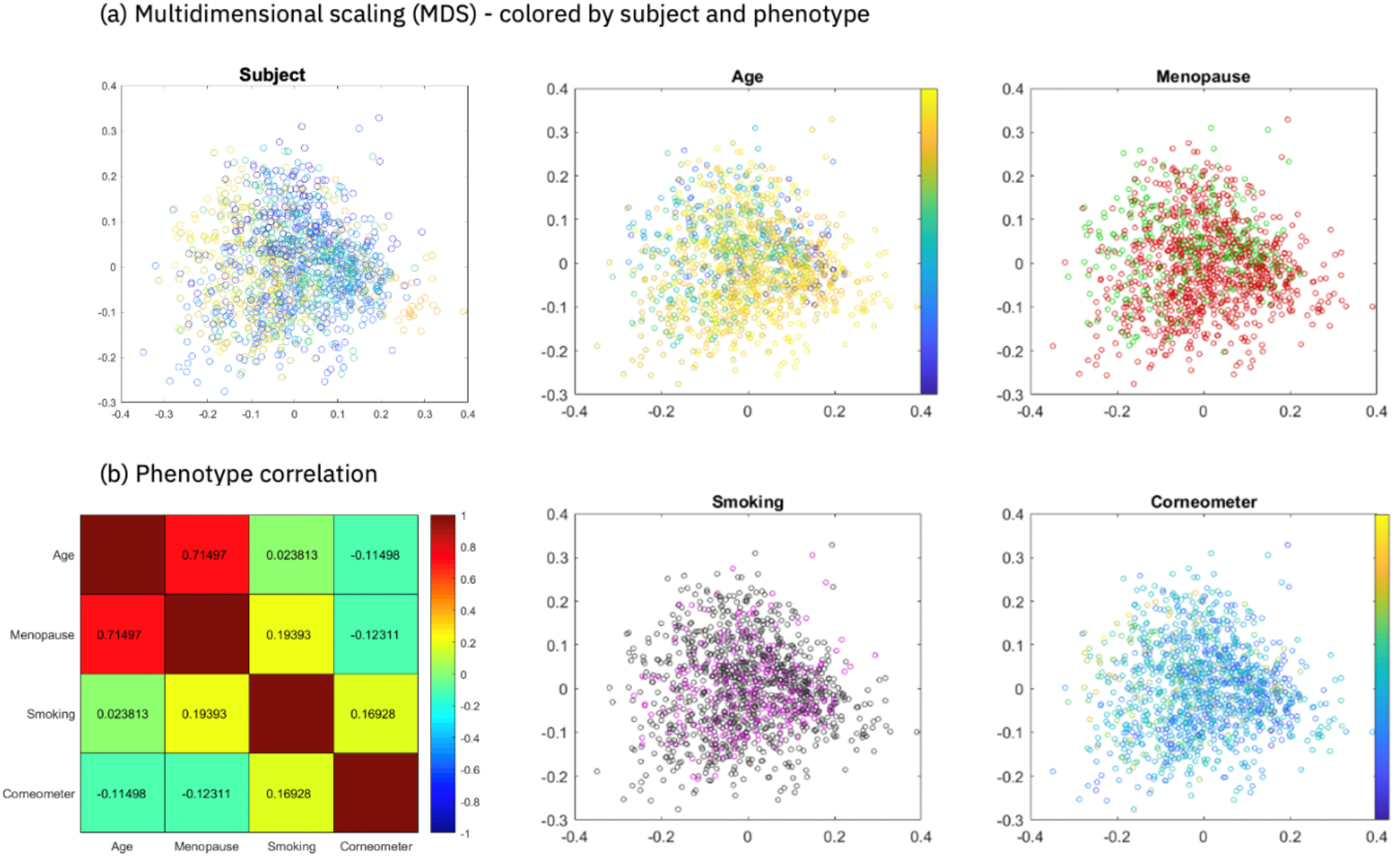
Phenotype prediction for Canada cohort. **(a)** Multidimensional scaling of pairwise sample Spearman distances. For subject, 63 randomly generated colours from blue to yellow are mapped to the 63 subjects. For age and corneometer, the colour scale is from blue to yellow, corresponding to the scale of smallest to largest observed phenotype value. For menopause red denotes “post-menopause”, and green “pre-menopause”. For smoking purple denotes “yes” and black “no **(b)** Pairwise Spearman’s correlation between phenotypes. Encoding for computing the correlation for the binary phenotypes: smoking no/yes=0/1, menopause pre/post=0/1.

Our EAI approach uses these features to predict skin hydration, age, menopausal and smoking status for the Canada cohort and examines the most impactful microbial genera when predicting the phenotype. We decided to investigate these phenotypes for the Canada cohort for the following reasons. Firstly, there is an increasing recognition of microbiome dysbiosis as a factor in atopic skin that is linked to psoriasis, eczema, acne, dermatitis and other disorders(25, 27– 29). Previous studies characterised microbiome signatures of different sites highlighting topographical and temporal variance in microbiome composition across dry, moist and sebaceous skin sites(26, 30–32). However, there is still a limited understanding of changes in the microbiome associated with cosmetic dry skin. We show that our EAI approach has the potential to provide insights on the effect of skin care and hygiene products on the molecular and microbiome composition of the skin (33). Moreover, being able to infer key changes in the skin microbiome associated with age may offer insights into the aging process, that could be used to develop products that counteract some effects of skin aging. Although a natural part of aging, the onset of menopause in women is a highly significant life event, especially for those women for whom the onset occurs earlier than expected. Menopause is currently diagnosed based on the symptoms, but a blood test to measure the hormone levels may be carried out for younger women. The power of predicting the onset of menopause through a simple scrub of the leg could be transformational to many women, as it points to the potential for a non-invasive diagnostic tool used as a condition monitor. Finally, previous studies have highlighted the strong impact of smoking on gastrointestinal microbiota(34),(35), and the differences in microbial community composition of the upper respiratory tract between healthy smokers and non-smokers(36). Although smoking is considered a contributing factor to skin aging and systemic health(37), its potential to influence the microbial community of skin has not been fully investigated.

Through the application of our carefully calibrated EAI approach we were able to accurately predict the four phenotypes and we identified skin microbial signatures associated with diverging values of each phenotype. To transform the biological insights provided by our EAI approach into actionable insights, we ranked the 186 genera by average SHAP impact and report the consensus top impactful genera for our bag of models. Our findings demonstrate the potential of explainable artificial intelligence in contributing to the growing body of knowledge of microbes and their host-interactions, and its future impact in research fields from cosmetic and medical research to forensic science and personal health monitoring(29, 33, 38, 39).

## Results

In the following sections we present the result obtained when predicting skin hydration, age, menopausal status and smoking status using the Canada cohort. To take into account possible confounding factors when predicting the four phenotypes for the Canada study, we checked if multidimensional projection of pairwise sample Spearman’s distances showed biases or separability of samples by subject and by phenotype (Figure 1(a)). The results indicate the samples are centred with no clear trends in either dimension of the two-dimensional scaling and they are not separable by subject and by phenotype. Moreover, we examined the connections between the skin hydration (corneometer), age, menopause and smoking phenotypes by computing their pairwise correlations (Figure 1(b)). Only age and menopause were found to be correlated (∼0.71 Spearman’s correlation), as expected. To address this, we considered confounding factors when predicting age and menopause and showed that we were able to predict age within the pre- and post-menopausal groups separately. Also, we highlighted the differences in predictive genera between the age and menopause models. With these checks in place, we proceeded with phenotype prediction. Table 2 shows a summary of predictive performance of our bag of models trained on the Canada cohort and evaluated on test set and 5-fold cross validation (CV) for each phenotype.

**Table 2.** Summary of predictive performance results of our fine-tuned bag of models trained on the Canada cohort and evaluated on a) test set, b) 5-fold cross validation (CV). For regression tasks (corneometer and age prediction) we show the average mean absolute error on test set (Test MAE) over the three ML model and the average MAE and standard deviation on 5-fold CV. Similarly, for classification tasks (menopausal and smoking status prediction) we report the average F1-score.

| Regression task | Test MAE | Ave MAE 5-CV | Std MAE 5-CV |
| --- | --- | --- | --- |
| Corneometer | 5.59 | 5.51 | 0.4 |
| Age | 4.51 | 3.97 | 0.39 |
| Classification task | Test F1-score | Ave F1-score 5-CV | Std F1-score 5-CV |
| Menopausal status | 0.92 | 0.93 | 0.020 |
| Smoking status | 0.93 | 0.91 | 0.026 |

### Explainable skin hydration prediction model generalises across cohorts

We investigated the prediction of multiple individual and combined measures of skin hydration, and found that corneometer, a widely used and robust method for assessing skin hydration(39), was the most appropriate measure for the Canada study (see section Phenotype analysis in Supplementary Information and Supplementary Figures 2-4). Efforts to enrich the corneometer score with other measures did not provide sufficient additional improvement in performance as to merit their inclusion. Higher corneometer score values correspond to higher skin hydration, thus we adopt the terms corneometer score and skin hydration interchangeably.

**Figure 2.**
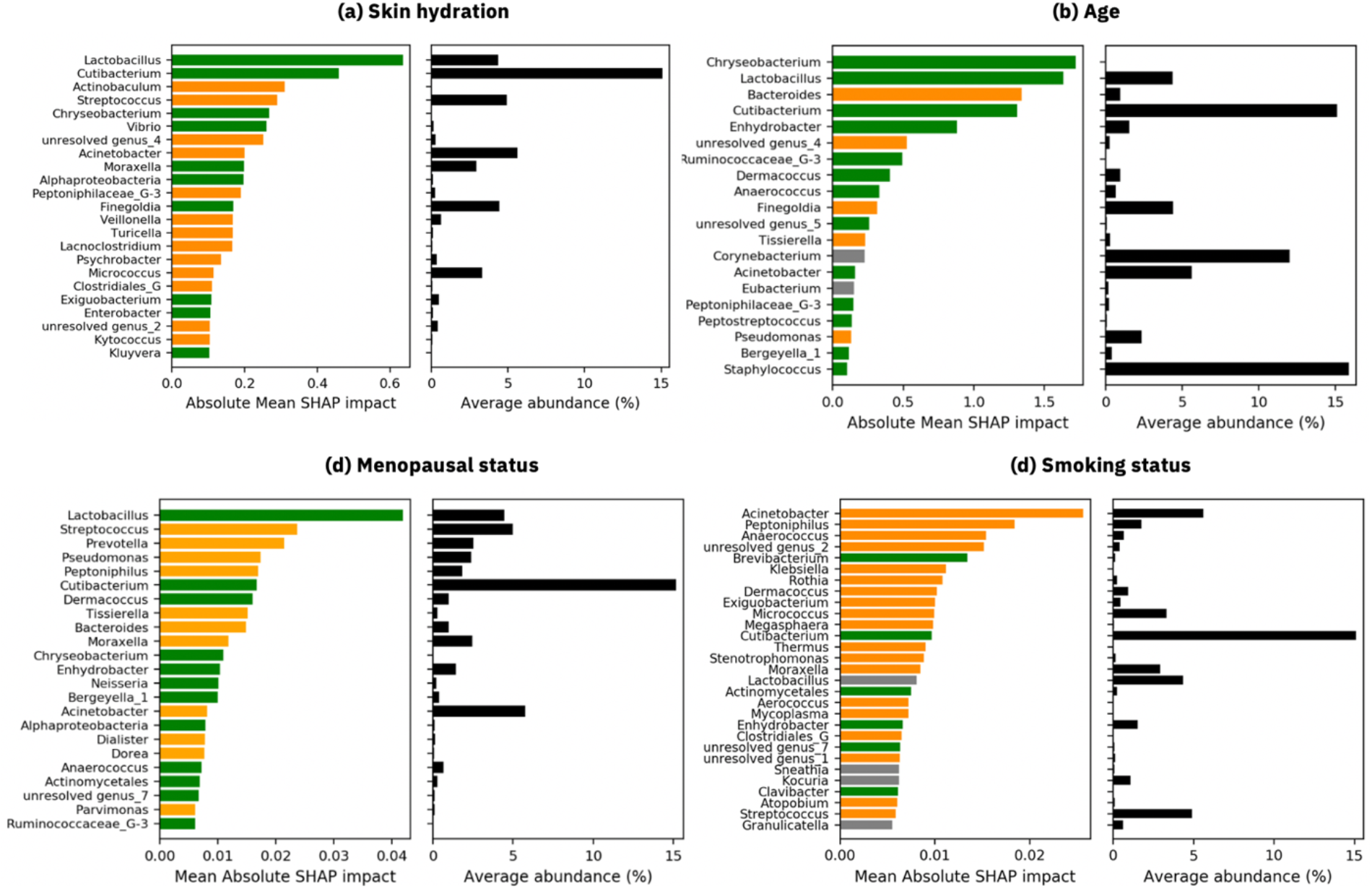
Most impactful genera when predicting the phenotype values with our EAI approach. **(a)** The top 23 genera are ranked by the absolute SHAP values when predicting corneometer, averaged across exemplars and the three different ML models. **(b)** The top 20 genera are ranked by the absolute SHAP values when predicting age. **(c)** The top 23 genera that have highest impact when classifying the set of exemplars using in two classes: pre-menopause and post-menopause. **(d)** The top 29 impactful genera when classifying the set of exemplars in two classes: smoking and non-smoking. On the left of each plot, green corresponds to the impact that the genus has, when more abundant in the microbiome, in predicting high skin hydration, young age, the class pre-menopause, or the class non-smoker, respectively. Orange corresponds to the impact that the genus has, when more abundant in the microbiome, in predicting lower skin hydration, older age, the class post-menopause, or the class smoker, respectively. Grey denotes genera for which the direction of impact was not clear, i.e. we did not find an agreement across the three ML models. On the right of each sub-figure (a-d), the average abundance of each genus across the entire set of samples is shown.

When examining the corneometer score vs. the skin microbiome, all the optimized machine learning models in our bag successfully predicted skin hydration levels from the measured microbiome (see Supplementary Figure 5). The highest Pearson’s correlation between predicted and true values, 0.71, was obtained with RF. However, XGBoost (∼0.65 Pearson’s correlation) and NN (∼0.53) were better at modelling real distribution of true corneometer values (see Supplementary Figure 5 and 7). Moreover, Table 2 and Supplementary Figure 6(a) reports the mean absolute error (MAE) on test and 5-fold CV. Results from cross-validation show that our models are stable and perform well on unseen data, when predicting corneometer.

The explanations for the corneometer predictions were provided in terms of impactful genera as computed by SHAP. A positive SHAP impact for a genus means that when the genus is more abundant in the microbiome sample, it contributes to an increase in the predicted score, while a negative SHAP value means the genus contributes to a decrease in the predicted score when more abundant. As the number of the most relevant features cannot be known *a priori*, we utilised the kneed algorithm(40) to discover plateauing of the SHAP explanation impact curve, indicating a cut-off for the number of features with the highest impact. In this case, the plateau was observed at 23 features. Figure 2(a) summarizes the most significant biological insights obtained while explaining the predictions of corneometer values. We found that 10 genera, when more abundant in the samples, are responsible for predicting higher skin hydration, while 13 genera are responsible for predicting lower skin hydration (see Figure 2(a)). The right-hand side of Figure 2(a) shows the average abundance of the impactful genera across the entire set of samples. Note that genera with high SHAP impact differ substantially in terms of their relative abundance in the source microbiome data (see for instance *Cutibacterium* and *Chryseobacterium* in Figure 2(a)). We observed a similar trend of most impactful genera not necessarily being the most abundant across all the samples when predicting the other three phenotypes, as presented in the following sections. The genera most determinant of better skin hydration were *Lactobacillus*, and *Cutibacterium* (formerly *Propionibacterium*) both of which have previously been reported as being associated with skin hydration(23, 24), whereas the genera *Actinobaculum, Streptococcus, Acinetobacter, Turicella and Micrococcus* were associated with decreased skin hydration. As the insights shown in Figure 2(a) can be extended to include the entire set of genera, our streamlined approach offers an opportunity to investigate how skin genera impact skin health.

To investigate the generalizability of our approach we extended the skin hydration analysis to include an UK study composed of 278 samples from 102 subjects and the same set of 186 OTUs as for the Canada study. The aim is to validate the skin hydration model and investigate how the model generalises across cohorts, while highlighting differences and similarities in the explainability results. We observed the error more than doubled when training and testing the bag of models only on UK samples, compared to when training and testing using only the Canada samples (e.g., 11.85 vs 5.33 MAE with XGBoost in Figure 3). Note that corneometer assumes values in a range between [8.1,78.4] with median ∼27.3 for UK cohort, while the corneometer range for Canada cohort is [2.4, 57.2] with median ∼24.73 (see Table 1 and Supplementary Figure 1 for more detail on the distributions). As such, the dissimilarity in predictive performance might be due to a) different distributions of corneometer values between the Canada and UK cohorts, b) the disparity in dimensionality of the two datasets (1234 samples x 186 OTUs vs 273 samples x 186 OTUs. In fact, when combining Canada and UK samples together for both training and testing the models, we obtained a MAE closer to the one using the Canada study only, respectively 6.37 and 5.33 MAE for XGBoost in Figure 3. Additionally, Supplementary Figure 8 reports MAE on 5-fold CV using Canada study only, UK study only and Canada and UK merged together. In addition, when using XGBoost trained on Canada samples to predict unseen UK samples, we obtained a MAE of ∼12.59, which is close to the 11.95 MAE obtained using XGBoost trained on UK samples to predict unseen UK samples (Figure 3). This result suggests that the models trained on Canada samples can be used to predict samples from the UK study obtaining comparable predictive performances.

**Figure 3.**
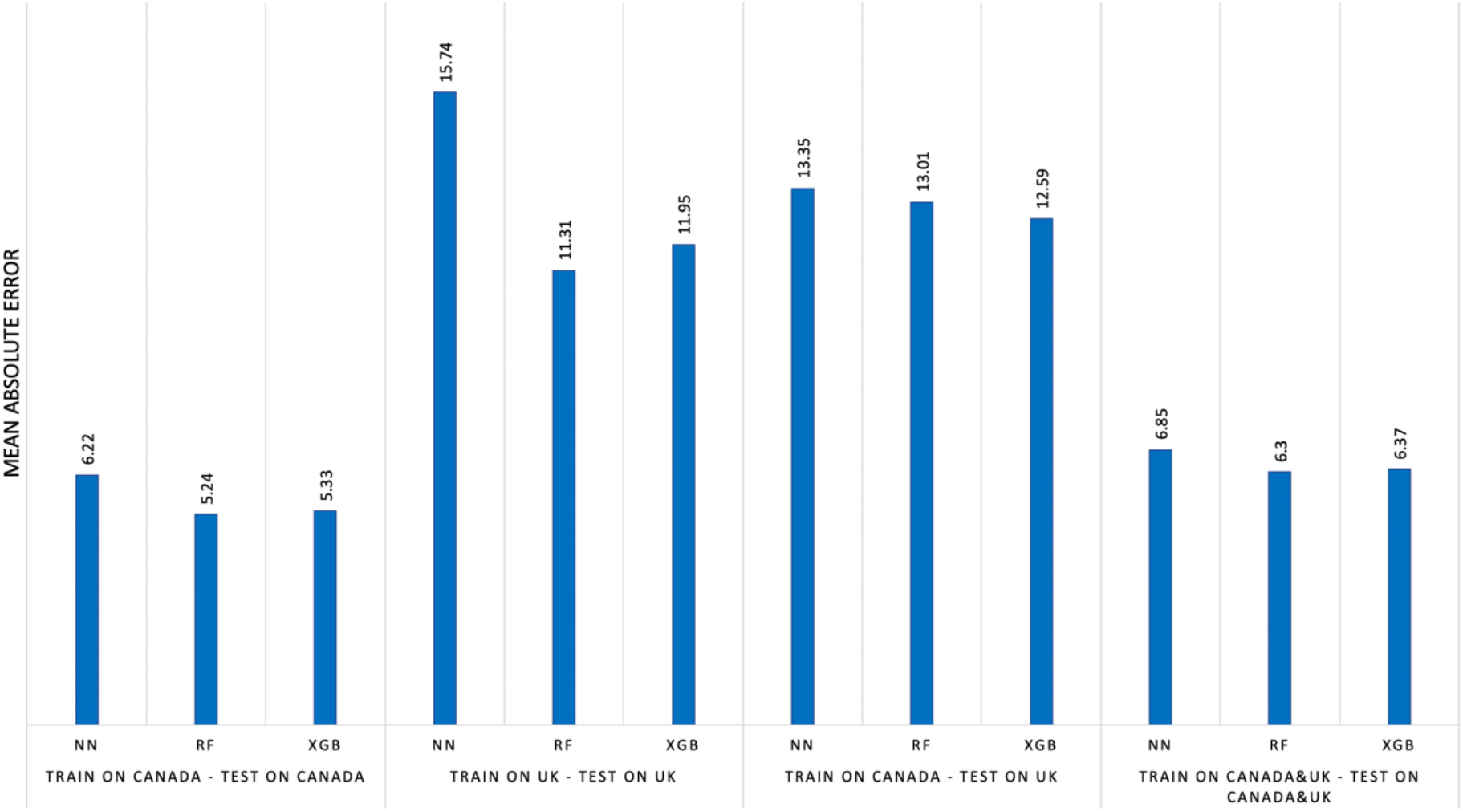
Predicting skin hydration (corneometer) using Canada and UK cohorts. Comparison of the mean absolute error (MAE) when – from left to right - 1) predicting corneometer for 20% of Canada samples with the model trained on 80% of Canada samples, 2) predicting corneometer on 20% of UK samples with the model trained on 80% of UK samples, 3) predicting corneometer for UK samples with the model trained on Canada samples, 4) predicting corneometer for 20% of the union of Canada and UK samples with a model trained on 80% of the union of Canada and UK samples.

Finally, we applied our EAI approach to explore the commonality and difference between the impactful genera for predicting skin hydration for the Canada cohort and the UK cohort separately. We found common impactful genera in both UK and Canada models. *Lactobacillus* and *Cutibacterium* (formerly *Propionibacterium*) are among the top 20 impactful genera driving the prediction of more hydrated skin (see Supplementary Figure 9) both of which have previously been reported as being associated with skin hydration(23, 24). *Micrococcus* and *Actinobaculum* are another example of top impactful genera driving the prediction of less hydrated skin for both UK and Canada models. In addition, there are some genera in the UK model, such as *Kocuria, Gordonia, Staphylococcus*, that do not appear in the list of the top impactful genera for the Canada model and vice versa (see Supplementary Figure 9 and Figure 2(a)).

### Accurate model identifies key microbes predictive of age

Following a recent study that found the skin microbiome of the hand and forehead to be the best in predicting age compared to the gut and the oral microbiome yielding predictions within 4 years of chronological age (42), we applied our EAI approach to predict age for the Canada cohort. Our EAI approach was able to predict the subjects’ age from the same set of skin microbiome samples achieving an average error within 4 years of chronological age, similarly to the error obtained in (42). Supplementary Figure 10 and 11 show the predictive performances of the three models on the test set. For example, with XGBoost we obtained a high Pearson’s correlation (*r* ≈ 0.88) between predicted and true values of age (Figure 10). Table 2 and Supplementary Figure 6(b) show the average MAE obtained by our bag of models on test set and 5-fold CV, while Supplementary Figure 14 shows the results on cross validation by subject and cross validation using 1 sample per subject (see Methods). CV results give an indication of how well the bag of models predicts age on unseen data from the Canada cohort.

Analogously to the skin hydration model, we applied the SHAP explainability analysis to understand the link between microbial communities on skin and age. Our EAI approach revealed that higher abundance of 13 genera drives the prediction of lower values of age, while higher abundance of 5 genera drives the prediction of higher values of age (see Figure 2(b)). In the age model, the knee point was observed at 20 features. For two of the genera, *Corynebacterium* and *Eubacterium*, among the top 20 most impactful we did not find a consistent direction of impact according to our bag of models. These features are shown as grey bars in Figure 2(b).

We found some overlap between genera identified as being important for both skin hydration and age models as well as differences in the impact of genera for the age model in comparison to the skin hydration model (Figures 2(a) and Figure 2(b)). The age model indicated a greater importance of *Chryseobacterium* and key genera, such as *Staphylococcus* and *Bergeyella_1*, that were not seen among the most impactful genera in the skin hydration model. This enabled the disambiguation of microbes whose effects mainly relate to either age or hydration. In fact, perhaps contrary to expected, skin hydration and age are not inexorably linked. This can be observed in Figure 1(a), which shows a very weak link between age and corneometer score (Spearman’s correlation score -∼-0.11). Our EAI approach indicated *Cutibacterium* as a key genus for predicting age. To further investigate this observation, we produced a stacked force plot (see Supplementary Figure 12), filtered on *Cutibacterium*. The figure indicates that high abundance of *Cutibacterium* seen in younger people falls dramatically after the age of 50; a similar trend has also been reported previously(43). This coincides with the well-known reduction in sebum after 50 years of age(44).

Finally, we used the UK cohort as a hold-out dataset to predict age with the bag of models trained on the Canada cohort. The MAE obtained predicting age for UK samples from models trained on Canada cohort is ∼9.63 years. The latter error is comparable to the average MAE (∼8.54 years) obtained when training our bag of models on UK samples to predict the age of unseen samples from the same UK cohort. As per the skin hydration model, the disparity in age prediction performance between UK and Canada cohorts may be due to the differences in true age distributions, dataset size, and country-specific differences in microbiome composition.

### Leg skin microbiome accurately predicts menopausal status – Canada cohort

To examine microbial changes associated with the menopause and investigate how these changes are also related to aging, we applied our EAI approach to classify samples of Canada cohort into pre-menopausal and post-menopausal status. The dataset includes 324 samples in pre-menopausal status and 876 in post-menopausal status. Only one subject, for a total of 34 samples, was in peri-menopausal status therefore was excluded from pre/post-menopausal prediction. The best F1-score of 0.93 on the test set was achieved by XGBoost (see Supplementary Figures 6(c)). The F1-score averaged over our three models is 0.92 (Table 2). Supplementary Figure 13 displays the confusion matrices for the three models, demonstrating how each model is able to accurately predict each class. In addition, Table 2 and Figure 6(c) summarise the performance results on 5-fold CV, while Supplementary Figure 14 shows the results on cross validation by subject and cross validation using 1 sample per subject (see Methods). These results demonstrate that menopausal status prediction is robust for different unseen test sets from the Canada cohort.

Our EAI approach revealed that higher abundance of 12 genera drives the prediction of the class pre-menopause, while 11 genera were identified as most impactful in predicting post-menopause when higher in abundance (Figure 2(c)). As menopause and age are strongly correlated, as expected, we investigated similarities and differences between the two models taking into possible confounding factors when predicting age or menopause. The model considered important a number of genera (*Lactobacillus, Cutibacterium, Enhydrobacter, Dermacoccus, Enhydrobacter*) already seen as key indicators in the age model, along with a key indicator of skin hydration (*Lactobacillus*). This is perhaps not surprising as menopause usually occurs in middle age and the skin is significantly affected by the aging process and menopause(45). Genera such as *Peptoniphilaceae_G-3 Finegoldia, Ruminococcaceae_G-3, Peptostreptococcus, Corynebacterium* appear among the most impactful for predicting age, while they do not appear in the menopausal model. On the other hand, *Actinomycetales, Moraxella, Acinetobacter, Dorea, etc*. are considered impactful for predicting menopausal classes and do not appear in the ranked list of most impactful genera of the age model. *Acinetobacter* is important in the age model to predict younger age, while it predicts post-menopause when higher in abundance in the menopausal model. It is also important to point out that the age range of women in the cohort in the pre- and post-menopausal status overlaps significantly, from 21 to 55 years old and 21 to 65 years old respectively.

We then trained our ML models for predicting age separately for pre-menopause and post-menopause samples. We were able to accurately predict age for the two sub-groups, showing that age prediction is robust (See Supplementary Figures 15-16). The latter results, together with the overlap in age span between pre and post menopause classes, and the differences observed in the ranked lists of impactful genera for the two models, indicate that the age and menopausal models differ from each other and may offer insights specific to the phenotypes. Furthermore, we examined the local explanation of a 41-year-old subject, who was correctly classified in post-menopausal status. This classification was made correctly showing that our model is not confounded by age since according to age the likely classification of the subject would be as pre-menopausal, with the majority of women reaching the menopause between the ages of 45 and 55. We found that the presence of genera such as *Prevotella* and *Acinetobacter* in this sample drove the correct prediction of the class post-menopause, while lower abundance of *Cutibacterium* had a small negative impact in predicting post-menopausal status (see Supplementary Figure 17). This is consistent with the explanations in Figure 2(c). Moreover, we found that higher abundance of genera that do not appear among the 23 most impactful for menopausal classification (i.e., *Micrococcus, Gordonia* and *Brachybacterium*) drove the correct prediction of post-menopause for this sample. This suggests it may be useful to extend the analysis to investigate local explanations of the predictions for specific samples of interest as well as the impact of the total set of genera in predicting the menopause.

### Key microbes on the leg discriminate smokers accurately - Canada cohort

With the aim of investigating microbial differences on the skin associated with smoking, we applied our EAI approach to predict smoker vs non-smoker subjects from the leg skin microbiome of Canada cohort. The dataset includes 286 samples from smokers and 948 samples from non-smokers. We were able to accurately predict the two classes with the all three ML models reaching an average F1-score over the three models of 0.93 (see Table 2). Supplementary Figure 14 shows the confusion matrices for the three ML models. Figure 6(d) and Table 2 show the performance results over 5-fold CV, while Supplementary Figure 13 shows the results of cross validation results by subject and cross validation using 1 sample per subject (see Methods).

We found that predicting the class smoker is driven by higher abundance of 19 genera (including *Acinetobacter, Peptoniphilus, Anaerococcus, Klebsiella*), while 6 genera (including *Brevibacterium, Cutibacterium, Actinomycelates*) were the most impactful for predicting the class non-smoker (Figure 2(d)). The knee point was found at 29 genera.

Finally, we looked at the split of smoker vs non-smoker against age. The age of smokers varies from 21 to 65 years old, while the subject’s age of non-smokers varies from 40 to 64 years old. We were able to accurately predict age for the two sub-groups (smokers and non-smokers) separately, showing that age prediction is robust despite possible confounding factors like smoking (See Supplementary Figures 18 and 19).

## Discussion

To summarize the explainability results across phenotypes, we compiled Table 3 illustrating the overlapping and uniquely impactful skin microbes per phenotype for the Canada cohort. Our study indicated that some members of the microbiome play a general role in wellbeing as overlapping sets of genera were involved in consistently predicting skin hydration, age, smoking and menopausal status when more abundant in the microbiome (see agreement in the top half of Table 3). The genus at the top *Cutibacterium* is impactful for predicting all four phenotypes in a consistent direction when more abundant. *Lactobacillus* is impactful for predicting higher skin hydration, younger age and pre-menopausal status, while *Streptococcus* is impactful for predicting lower skin hydration, post-menopausal and smoking status. The following fourteen genera are impactful for predicting at least two phenotypes each. Nevertheless, accurate predictions of each phenotype required distinct combinations of impactful genera (see differences in the bottom half of table of Table 3). For example, abundance of *Turicella* and *Veillonella* are uniquely driving the prediction of lower skin hydration, while *Staphylococcus* and *Coryneobacterium* are most impactful only in the age prediction model. Higher abundance of *Neisseria* uniquely drives the prediction of post-menopausal status, while higher abundance of *Rothia* identifies smokers. *Peptoniphilacae* is an example of a genus having opposite impacts on two phenotypes (lower skin hydration and younger age).

**Table 3.**
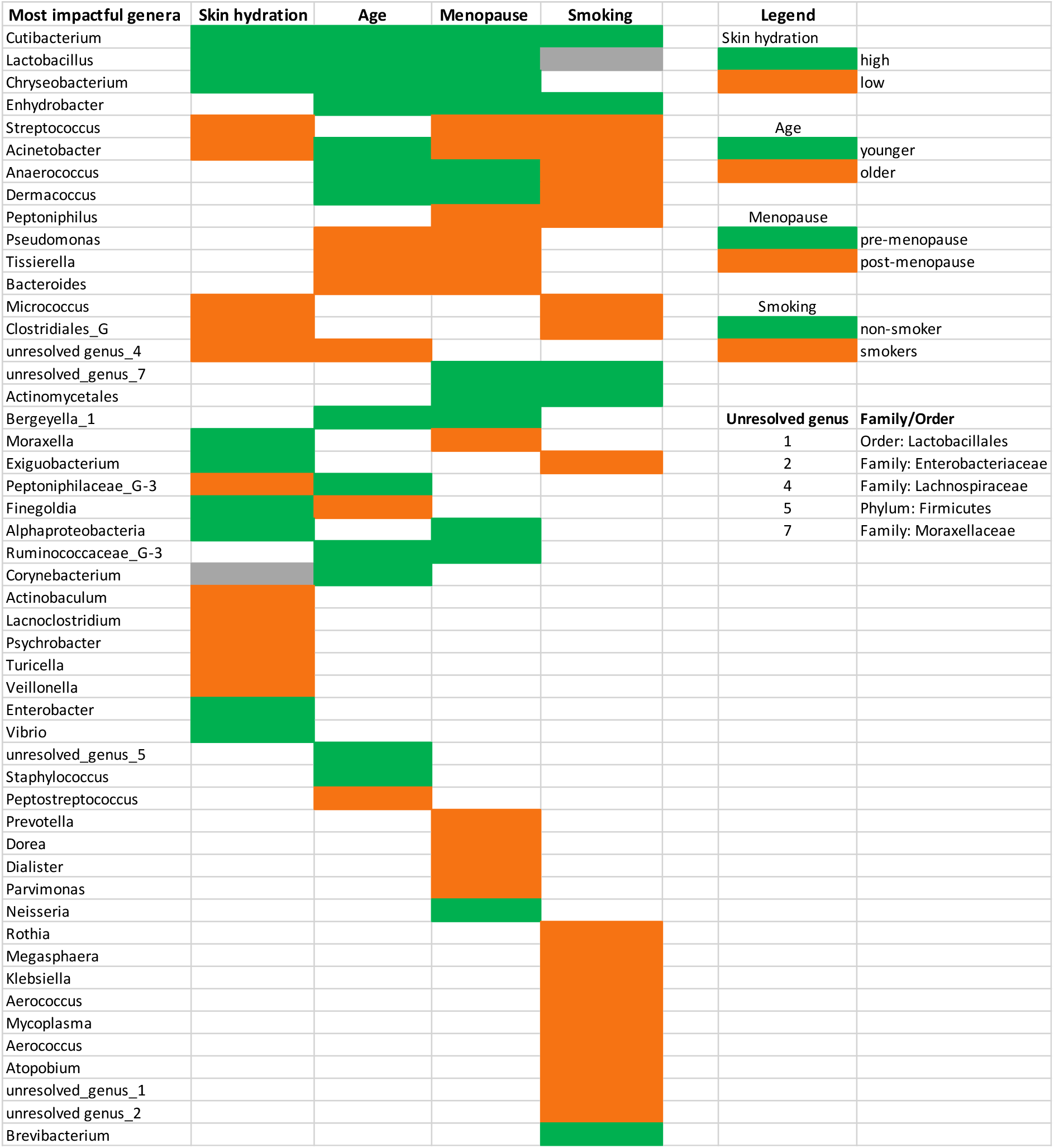
Summary of insights inferred from our EAI framework when predicting skin hydration, age, menopausal status, and smoking status for Canada cohort. The table shows the union of the most impactful genera for each phenotype. The colors reflect the impact that each genus has in predicting a particular phenotype when higher in abundance. Information about the order or family of the unresolved genera is also provided. For example, abundance of Turicella and Veillonella are uniquely driving the prediction of lower skin hydration, while Megasphaera and Klebsiella are most impactful only in the age prediction model. Note that we did not include Eubacterium in this table as its direction is not clear for age prediction and it does not appear in the list of most impactful features for the other phenotypes. Similarly, we did not include Sneatia, Kocuria, Granulicatella as their direction for the smoking model is not clear and they did not appear among the top impactful features for the other phenotypes.

We visualized the average relative abundance per subject of the impactful genera for each phenotype (Figure 4(a)). By comparing the microbial community profiles of the subjects, it is not discernible how the impactful genera differentiate between phenotype groups or values. Hence, our explainable artificial intelligence approach has the potential to uncover the subtle interplay of signals present in the microbiome that is far from obvious when viewing the input data.

**Figure 4.**
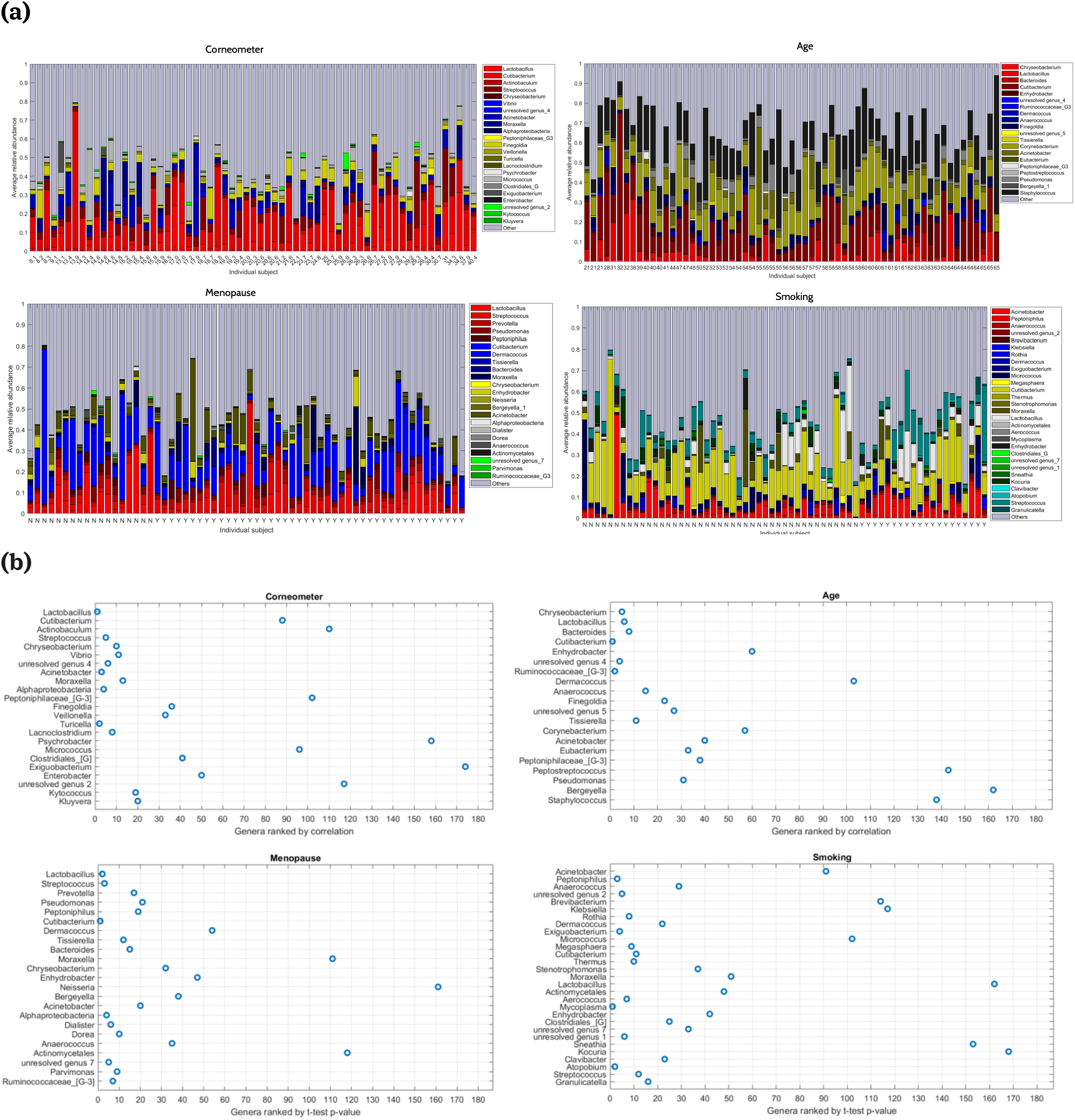
**(a)** Visualizing the relative abundance of impactful genera when predicting skin hydration, age, menopausal and smoking status for Canada cohort. The subjects are ordered by phenotype values (x-axis). For each subject we averaged the relative abundances of each genus across all samples for that subject, after removing any outlier samples that had correlation < 0.5 with more than half of the samples for that subject. The impactful genera per phenotype are then visualized, and all other genera labelled as “Others”. **(b)** Visualization of top impactful genera per phenotype from ML (vertical axis, most impactful genus on top) versus their rankings in independent statistical tests (horizontal axis, the total number of genera is 186).

**Figure 5.**
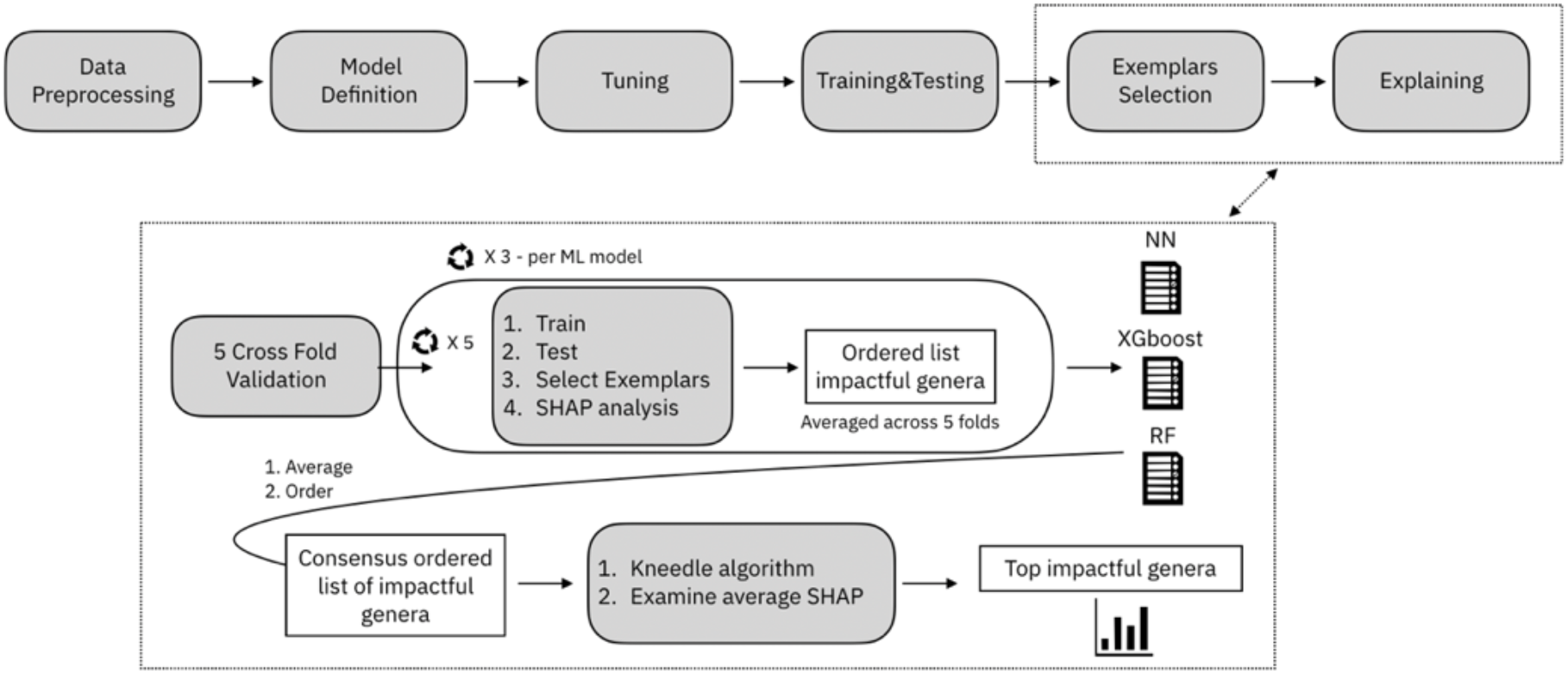
Outline of our EAI approach.

Additionally, we compared the impactful genera from machine learning to the results from statistical tests that independently associate each genus with the phenotype (see Figure 4(b)). We observed overlap between the top genera from ML and from statistical testing, however, there were notable exceptions also. For corneometer the 2^nd^ most impactful genus *Cutibacterium*, was only ranked 90^th^ when the ML sensibly links it with skin hydration due to its association with acne and propensity for wet skin. The 3^rd^ most impactful genus *Actinobaculum*, that has previously been associated with skin infection but is typically observed at low abundance(46), was ranked 110^th^ by correlation. For age *Dermacoccus, Peptostreptococcus, Staphilococcus, Bergeylla* are among the top impactful genera in the age model but only ranked below 100. Similarly, *Moraxella, Neisseria, Actinomycetales* are among the most impactful genera in the menopausal status model but ranked below 100 in the t-test. For smoking, the top ML feature *Acinetobacter* was only at rank 91^th^ according to the t-test and other 6 genera were ranked below 100. Species within the *Acinetobacter* genus have been linked with respiratory and bloodstream infections which could explain their usefulness in smoking prediction with this being reflected in the leg microbiome(47). This indicates the set of most predictive genera inferred by our EAI approach can differ from the ones highlighted by a standard statistical approach that consider each genus independently. As such, our approach can be used in addition to a standard approach to investigate predictive genera that could be missed if at the bottom of the ranked list produced by a statistical test.

Our findings on the predictive power of the leg skin microbiome point to the possibility of using more accessible microbiome samples to investigate phenotypes (e.g. smoking) that are not believed to be directly associated with the microbiome of the sampled body site. The increase of *Actinobacteria* previously observed in the gut microbiome of non-smokers is unexpectedly in agreement with the fact that we found *Brevibacterium, Cutibacterium, Actinomycelates, Clavibacter*, all belonging to the phylum *Actinobacteria*, drive the prediction of non-smokers (see Figure 2(d)). *Streptococcus, Actinomyces, Megasphaera, Atopobium* were previously identified as indicators of smokers in the respiratory tract(36), which is also in agreement with our findings. Increased levels of *Megasphaera* have been found in the oral cavity of smokers(48), with *Megasphaera* being among the impactful genera for predicting smokers in Figure 2(d). Smoking related increases in the anaerobic *Actinobacteria* OTUs from *Actinomyces* spp., *Rothia mucilaginosa* and *Atopobium* spp. were found in the oral microbiome(49). Similarly, our EAI approach found that *Rothia* and *Atopobium* predict smokers when more abundant in the microbiome samples. Cigarettes themselves harbour a broad range of potential pathogens, including *Acinetobacter, Clostridium*, and *Klebsiella*, that have been found to be predictors of smokers when abundant in the leg skin microbiome (see Figure 2(d)). The similarity between the genera that discriminate smokers from non-smokers in our skin model and the genera identified as discriminators of smokers in previous studies of the oral and gut microbiome is encouraging. Despite the fact that smoke is not in direct contact with skin on the leg, we found the leg skin microbiome changes in response to smoking, and in a manner that is consistent with previous findings from different body sites. This further confirms systematic changes occurring across the human body in cigarette smokers. It also suggested the microbiome of distant body sites may be affected by smoking, hence pointing to the fact that extremely complex interactions between body locations or tissues are yet to be examined.

Likewise, microbiome sequencing from vaginal swab samples found a similar composition of major vaginal bacteria at the genus level in pre- and post-menopausal women but with altered proportion(50). The previous study observed high levels of *Lactobacillus* (64.4%) in pre-menopausal women that were lower post-menopause (24.4%), replaced by *Prevotella* (11.4%) and *Streptococcus* (5.1%) among others. This reflects the observations here where *Lactobacillus* is most impactful when more abundant for predicting pre-menopause while *Prevotella* and *Streptococcus* are most impactful when more abundant for predicting post-menopause (see Figure 2(c)). Here the skin microbiome appears to imitate the vaginal microbiome closely which could explain its ability to explain menopausal status.

We showed that genera with a relatively high SHAP impact can differ substantially in terms of their relative abundance in the source microbiome data. Therefore, our EAI approach identifies the most impactful genera across the samples even when these are not the most abundant genera. For instance, although *Chryseobacterium* is one of the least abundant genera across the entire set of samples, it is the most impactful in the age prediction model. We propose that all genera with a relatively high SHAP impact are likely to either play a biologically causative role or offer value as a non-causative indicator organism and thus are worthy of additional study.

We took into account possible confounding factors when predicting the four phenotypes for the Canada study. Of the four phenotypes, only age and menopause were found to be correlated (∼0.71 Spearman’s correlation, Figure 1(b)). To address this, we confirmed that we were able to accurately predict age separately for the pre and post-menopausal groups (See Supplementary Figures 15 and 16). Through our explainability analysis we demonstrated how the age and menopausal models share common impactful genera while still differing from each other. Therefore, although the phenotypes are correlated, the age and menopausal models may offer insights specific to each phenotype. Also, we were able to accurately predict age separately for the sub-groups (See Supplementary Figures 17-18), demonstrating that age prediction is robust despite possible confounding factors like smoking and menopausal status. Additionally, the microbiome profiles of the samples are not separable by phenotype by applying multidimensional projection (Figure 1(a)), yet our EAI approach is able to accurately predict the phenotype values.

Furthermore, we evaluated our models by performing 5-fold CV on both the Canada and UK cohorts. The results of cross validation, summarised in Table 2 and Supplementary Figure 6, show that our models perform well on different unseen test sets.

For the Canada cohort we performed three additional validation steps. Firstly, through multidimensional projection of pairwise sample Spearman’s distances we showed the samples are centred with no clear trends in either dimension and they are not clustered by subject (Figure 1(a)). Secondly, we performed cross validation by subject, e.g. all samples of one subject were set aside for testing while the models were trained on the remaining subjects’ samples (see Methods). In this case for regression tasks, we observed a small increase in average MAE compared to 5-fold CV on the entire set of samples (Supplementary Figure 14(a)). For example, for corneometer prediction we obtained an average MAE of ∼5.51 on 5-fold CV and an average MAE of ∼7.53 on CV by subject. For classification tasks (smoking and menopausal status prediction) we observed a small decrease in average accuracy (Supplementary Figure 14(a)). For smoking status prediction, we observe a small decrease in average F1-score (∼0.90) across the three models when performing cross validation by subject compared to the average F1-score from 5-fold CV (∼0.91). Note that the decrease in performance may be due to the difference in number of folds (5 vs 63) and the difference in number of samples per fold (∼246 samples per fold for 5-fold CV; see distribution of samples per subject in Supplementary Figure 1). Thirdly, we investigated if it was possible to predict the four phenotypes using only one sample per subject without suffering a drastic decrease in predictive performance. After removing the one subject in peri-menopausal status, we chose to keep the first sample taken from each subject, i.e., at the earliest date, therefore picking 62 samples in total. We then trained and tuned our models to predict the four phenotypes on these selected samples only. Supplementary Figure 14(b) reports the performance results of 5-CV using RF and XGBoost on the 62-sample subset. The results of cross validation using 1 sample per subject are comparable to the results of cross validation by subject (Supplementary Figure 14a and 14b). As such, we showed that using only 62 samples of 62 different subjects does not drastically decrease the predictive performance of our models. On the contrary, the small increase in error and small decrease in accuracy are expected and due to the much smaller sample size (62 vs 1234 samples) used to train our models. All the validation results above demonstrate that our ML models are robust in predicting the four phenotypes as they show comparable predictive performances when trained on different subsets of samples and tested on different unseen test tests.

In addition, we demonstrated the generalisability of the skin hydration and age models to a different cohort (UK). We showed that we can use our bag of models trained on Canada samples to make predictions on UK samples. The error obtained in the latter case was in fact comparable to the error obtained training our bag of models on UK samples only. Moreover, we found common impactful genera in both UK and Canada models. *Lactobacillus* and *Cutibacterium* (formerly *Propionibacterium*) are among the top 20 impactful genera driving the prediction of more hydrated skin. *Micrococcus* and *Actinobaculum* are another example of top impactful genera driving the prediction of less hydrated skin for both UK and Canada models.

Finally, while predictive performance is important, the focus of this work is our explainable AI approach. Providing model explanations can help us to understand the link between microbiome composition and phenotypes, and also importantly build trust in the model predictions.

In this study we investigated microbial taxa at genus level, however our EAI approach can be similarly applied to the investigation of changes at species or strain level. Moreover, while we performed a particular analysis with one bioinformatics pipeline (to identify genera and abundances from bacterial 16S rRNA gene sequencing), this approach extends to exploring how microbial features from shotgun metagenomic sequencing are impactful in predicting phenotypes. This would involve taking into account the entire genetic material of a microbial community (including fungi and viruses), providing insights related to genes and their associated biological functions. In general, the ability to predict specific phenotypes from non-invasive microbiome sampling can perform a similar revolutionary role as the collection and sequencing of human DNA from cheek swabs, which has powered the gathering of massive data sets and thus the genomic revolution in personalized medicine.

## Conclusion

We developed an explainable AI (EAI) approach to predict phenotypes and explain the predictions in terms of changes in microbiome composition related to diverging phenotype values. The results presented in this study demonstrate the power of explainable AI in accurately predicting diverse phenotypes such as skin hydration, age and, surprisingly, menopausal and smoking status from the leg skin microbiome and in inferring microbial signatures associated with each phenotype. In addition, we investigated the application of the skin hydration model to a second cohort and compared predictive performance and explainability.

Our study focused on personalised care and wellbeing. However, it is straightforward to appreciate how this approach is broadly applicable in healthcare, representing an advancement in the deployment of microbiome-based and non-invasive diagnostics for practitioners (e.g. dermatologists) and in the design of personalised treatments. Impactful future work will likely involve additional independent studies and analysis at species or strain level to investigate health-related phenotypes and how the explanations generalise across more populations.

Our EAI approach will enable the community to expand and build on this initial work as the explanations may offer new insights into the complex interactions between microbes and their host, possibly leading to new interventions to adjust the microbiome for improved health outcomes.

## Methods

### Clinical design and data collection

#### Study participants

A total of 63 Canadian women between 21-65 years of age (average 52.71 years) took part in this double blind randomised, balanced application study, between April and July 2017. In addition, 102 women aged between 19-55 years (average 38.03 years) took part in a UK based study. Key inclusion criteria included subjects being in good general health, not being pregnant or lactating and having intact non-diseased skin with minimal hair (on test sites). Key exclusion criteria included pregnancy or lactating mothers and use of moisturisers on the proposed test site 3 or more times per week. A full list of the inclusion and exclusion criteria can be found in Supplementary Notes.

#### Study Design

The study was carried out by an independent third party. Study visits include visual and instrumental assessments, as well as buffer scrub sampling. Additionally, subjects were asked to confirm their age, smoking and menopausal status at the start of the study. There was no conditioning (wash-out) phase required for this study.

#### Assessment of skin condition

Visual assessments of leg skin include dryness and erythema grading. Visual dryness assessments were made by an expert assessor with dryness scored on comparable scales from 0-6 (Canada) and 0-4 (UK) both with ½ point increments allowed. The descriptors for 0-4 skin condition in terms of severity are similar across scales. Instrumental assessments (Skicon™, corneometer, and pH) were taken following the visual assessments and a total combined acclimation period of a minimum 25 minutes to an environmental controlled room. The Skicon-200EX (IBS Co./AcaDerm) measures skin hydration by skin surface conductance, and for the Canada study a Corneometer CM 825 Courage+Khazaka) was used to measures skin hydration by capacitance (the skin’s ability to conduct a small electrical current). For logistical reasons the UK study used a Courage and Khazaka Multi-Probe Adaptor Corneometer (MPA 6) for the same purpose both instruments are widely used for this purpose and offer comparable function, performance and sampling area. Skin surface pH in the Canada study was measured using a pH meter (Courage+Khazaka).

The corneometer measurement was the most amenable for machine learning models (see Phenotype analysis in Supplementary Notes), therefore we focused on it in this study. Between 3 and 5 corneometer readings per test site were obtained and the probe was moved slightly (overlapping may occur) with each reading, but still within the site. Pressure on the skin surface was measured by means of a probe spring and was 3.5 Newtons onto the area measured. The area of skin in contact with the probe was 49 mm^2^.

#### Microbiome sampling

After all instrumental measures, cup scrubs buffer washes were collected using the method of Williamson and Kligman(51). Using a sterile plastic disposable pipette 2.5 ml of buffer wash solution (PBS+0.1% Triton X-100) was aliquoted into the sampling ring and gently agitated for one minute with a sterile rod. The sampling fluid was collected using a sterile disposable pipette and placed into a sterile tube. The sampling procedure was repeated with another 2 ml aliquot of buffer wash material. After agitation, this aliquot was added to the first. Samples were frozen at −20 °C within 10 minutes until DNA extraction. Skin microbiome samples were taken by the cup scrub method at four different points, with either one or two samples being taken on each leg, as described above. Samples were taken from the same individual at different time points giving an overall total of 1234 microbial samples from the 63 subjects.

#### DNA extraction

DNA extractions were carried out by QIAGEN (Germany). Frozen buffer scrub samples were allowed to thaw at room temperature and cell pellets sedimented by centrifugation at 13,000 rpm, 4^°^C for 10 minutes. Following removal of the supernatant the cell pellet was resuspended in 500µl TE buffer and transferred to a 96-well Lysing Matrix Plate B (MP Biomedicals, USA). Addition of 3µl of Ready-Lyse lysozyme (Epicentre, 250U/µl) was followed by incubation with agitation at 300rpm, 37^°^C for 18 hours. Following lysis DNA was extracted and purified using the QIAmp UCP DNA Micro Kit (Qiagen, Germany) following the manufacturer’s instructions. DNA was eluted into AUE buffer and frozen until processing for sequencing library preparation.

#### NGS library preparation and sequencing

Oligonucleotide primers targeting the V1-V2 hypervariable region of the 16S rRNA gene were selected. PCR primers (details in Supplementary Information) were a modified version of the standard 28F and 338R primers which contain additional recognition sequences to facilitate nested PCR to add Illumina sequencing adapters and index sequences to resulting amplicons. PCRs consisted of 0.25 μl (10 μM) of each primer, 10 μl of HotStar Taq Plus Mastermix (Qiagen), 5 μl of template DNA and 4.5 μl molecular grade water (Ambion, Thermofisher). Samples were amplified using the following parameters: 95°C for 5 minutes, then 10 cycles of: 94°C for 45 seconds, 65°C for 30 seconds, and 72°C for 60 seconds, with a final extension of 10 minutes at 72°C using a Dyad Thermocycler (MJ Research). PCR products were purified using Ampure SPRI Beads (Beckman Coulter, California, USA). A second round PCR incorporated Illumina adapters containing indexes (i5 and i7) for sample identification utilising eight forward primers and twelve reverse primers each of which contained a separate barcode allowing up to 96 different combinations. Second round PCRs consisted of 0.5 μl (10 μM) of each primer, 10 μl of 2 x Kapa Mastermix (Roche, Switzerland) and 9 μl of purified sample from the first PCR reaction. Samples were amplified using the following parameters: 98°C for 2 minutes, then 15 cycles of; 20 seconds at 95°C, 15 seconds at 65°C, 30 seconds at 70°C with a final extension of 5 minutes at 72°C. Samples were purified using AMPure SPRI Beads before being quantified using Qubit fluorimeter (Invitrogen, California, USA) and assessed using the Fragment Analyzer (Advanced Analytical Technologies, Iowa, USA). Resulting amplicon libraries were taken forward and pooled in equimolar amounts using the Qubit and Fragment Analyzer data and size selected on the Pippin prep (Sage Science, Massachusetts, USA) using a size range of 300–700 bps. The quantity and quality of each pool was assessed by Bioanalyzer (Agilent Technologies, California, USA) and subsequently by qPCR using the Illumina Library Quantification Kit (Kapa) on a Light Cycler LC480II according to manufacturer’s instructions (Roche, Switzerland). Each pool of libraries was sequenced on a flowcell of an Illumina HiSeq with 2 × 300 bp paired-end sequencing using v3 chemistry (Illumina, California, USA).

#### Informatics processing

The sequence data were processed as follows. Illumina adapters and PCR primers used for initial 16S rRNA gene amplification were removed from each fragment using Cutadapt version 1.14(52). Sickle version 1.33(53) was used to quality trim DNA reads using a minimum quality value of 28. Reads shorter than 100 bp following quality trimming were discarded. If a read was discarded during this process its corresponding pair was also discarded. Reads passing filtering were merged using PANDAseq(54) version 2.9 to generate overlapping contigs with a minimum overlap of 20 bp and a minimum amplicon length of 200 bp. The resulting overlapped reads were de-replicated using VSEARCH version v1.9.6(55) and searched against a BLAST database composed of the HOMD, HOMD extended and Greengenes sequences (HOMDEXTGG) described by Al-Hebshi *et al*(41). Genus level taxonomic classification was then performed for both datasets in a closed reference fashion against the same database HOMDEXTGG version 14.51, as previously described in (41), (56) at 99 % identity across 98% of the read length. Reads not classified by this process were discarded. Reads were marked *unresolved* when there was no agreement between the databases they were compared against. This process resulted in 1260 taxonomically classified operational taxonomic units (OTUs). The resulting classification table and associated representative sequences, selected as the most abundant sequence for each classified taxon, were used as inputs for QIIME(32)(57) (Quantitate Insights into Microbial Ecology) version 1.9.1.

The microbial sequence count table and metadata, in biom file format, was loaded into the calour library(58). The loading process filtered out samples with fewer than 1000 reads and then rescaled each sample to have its counts sum up to 1000 (by dividing each feature frequency by the total number of reads in the sample and multiplying by 1000). After loading, the data underwent two rounds of filtering and the remaining features were collapsed at the genus level. The first round of filtering removed low-abundance features (OTUs with total count less than 10 across all samples). The second filter removed OTUs with low prevalence (features occurring in <1% of the samples). The values for three bacterial genera that had been identified as contaminants, *Burkholderia, Dietzia, Mycobacterium*, were removed. After filtering and collapsing the OTUs at genus level we obtained a total of 186 OTUs and 1234 samples that were used in subsequent analyses.

#### Phenotype correlations, sample visualization, and feature ranking by statistical tests

Pairwise phenotype correlations were computed by Spearman’s correlation. Each subject only contributed once for each correlation measure, for corneometer and age the median of measured values per subject was used. To compute Spearman’s correlation for binary phenotypes (smoking and menopause status) we used the encoding smoking no/yes=0/1, menopause no/post=0/1. Pairwise sample distances were computed with Spearman distance between feature (genus) vectors per sample. Multidimensional scaling was applied to project the distances into two-dimensional sample representations for visualization. Statistical tests were applied to rank features as follows. For the binary phenotypes (smoking and menopause status), two-sample t-test was applied independently to each feature vector vs. the phenotype vector, and features were ordered by p-value from most to least significant. For age and corneometer, Spearman’s correlation between each feature and phenotype was applied, and features were ordered by absolute value of the correlation. When there were multiple measures of corneometer per subject, the median value was used.

### Explainable AI approach

The first step in our EAI approach, shown in Figure 5, consists of pre-processing and filtering the OTU table as described in the section above. Next, the ML models are defined depending on the prediction task: regression for predicting skin hydration and age; classification for predicting menopausal and smoking status. We split the dataset in two parts, 80% for training and 20% for testing. For each prediction task, the hyperparameters of each ML model are tuned performing 5-fold cross validation (CV) on the training dataset. Once the best hyper-parameters have been selected, each tuned model is trained and tested. The last two steps of our approach consist of selecting a subset of samples, *exemplars*, in the test set and applying explainability methods to interpret and explain the predictions made by the trained models on those exemplars.

#### Explainability of the predictions and exemplar selection process

Examining the ML models for explanations of their predictions is an important active field of research. As tree-based methods are more directly interpretable, recent research is focused on inspecting neural networks to try to ascertain the importance of each feature’s contribution to the prediction. Work such as DeepLIFT(59), DeepExplain(60), and SHapley Additive exPlanations (SHAP)(20) inspect the gradients of the models to determine the impact of different features. To investigate the mechanisms underlying the predictions, we used SHAP for its ability to work with any machine learning model, such as XGBoost and RF, not just neural networks. SHAP combines game theory with local explanation enabling accurate interpretations on how the model predicted a particular value for a given sample. The explanations are called *local explanations* and reveal subtle changes and interrelations that are otherwise missed when these differences are averaged out. Local explanations allow the inspection of samples that have extreme phenotypes values, e.g. very hydrated skin or very dry skin.

In order to provide robust explanations, from all the explanations provided by SHAP for the entire dataset, we selected explanations for *exemplar samples or examplars* i.e., samples in the test set for which the predictions lie very close to the actual measured values. More precisely, for regression tasks (corneometer and age prediction) we selected as exemplars the samples in the test dataset for which predictions were < 10% different from the actual values. For classification tasks (smoking and menopausal status prediction) we selected samples that were correctly classified. The exemplars were then examined using SHAP to assess how well the genera considered most impactful by the ML models matched existing biological knowledge. More precisely, we applied SHAP to the set of exemplars and obtained the list of features (microbial genera) in decreasing order of impact (i.e., SHAP absolute mean value), so that genera at the top were the most impactful in predicting the given phenotype.

As the set of exemplars differs depending on the chosen test set, we performed 5-fold CV for each ML model. For each genus, we computed the average SHAP impact over the 5 folds (corresponding to 5 different exemplar sets) and we re-ordered the list of genera based on their average SHAP impact in descending order. This gave us a robust ordered list of impactful genera for each model (RF, XGBoost and NN). Finally, we took the average SHAP impact of each genus over the three lists computed at the previous step and re-ordered once again all the genera based on their resulting impact. In this way we cross-validated SHAP results obtaining a single and robust consensus ranked list of impactful genera representing an agreement of the three models (see Figure 5).

Note that the average number of samples in the test set was 247 across 5-fold CV, while the average number of exemplars in the age model was 190 and 66 in the corneometer model. The average number of exemplars in the menopausal classification was 225, while the average number of exemplars in the smoking model was 223.

Finally, we applied the kneedle algorithm(40)(https://github.com/arvkevi/kneed – with the parameters curve=‘convex’, direction=‘decreasing’, interp_method=‘polynomial’) to find a plateau in the SHAP impact curve for the consensus list of impactful genera. We then examined the top *x* impactful features, where *x* is the knee point representing the beginning of the plateau. To assign a direction of impact for each genus, we computed the average SHAP impact (without taking its absolute value) of the top 20% of exemplar samples for which the genus was most abundant. For each genus and each ML model in our bag of models, we examined the sign of the SHAP impact averaged across 5 folds. A positive sign indicates that the genus had a positive impact in predicting a phenotype value when it was more abundant in the microbiome, while a negative sign indicated that the genus has a negative impact on the prediction when higher in abundance in the microbiome. We then assigned a direction to the impact of the genus under analysis if it was consistent across the three models. The results of this process for the four phenotypes are visualized in Figures 2(a-d). All the insights obtained across phenotypes are summarised in Table 3.

#### Machine learning models

We evaluated the application of three machine learning models - random forest (RF)(17), XGBoost(18) and neural network (NN) (19) - for predicting the four different phenotypes from microbial genus count tables. We developed our neural network using TensorFlow(61) to construct a multilayer perceptron (MLP). The MLP architecture, implemented in Keras, is shown in Supplementary Figure 20. We optimised the parameters of the neural network’s architecture using a Bayesian optimization algorithm(62). The input layer contains the same number of neurons as the number of features in the input data (186 neurons). The input layer is followed by a batch normalisation layer with rectifying linear unit (ReLU) activation function. This is followed by 4 blocks, where each block contains a batch normalisation layer, a fully connected layer with 100 neurons, followed by ReLU activation function. The penultimate layer is a dropout layer (we used 40% dropout). The final layer is either one neuron in the case of regression for corneometer and age prediction, or a softmax layer in the case of classification tasks, with as many neurons as there are classes. The model was trained in batches of 80 samples. To further aid generalization of the neural network we applied early stopping, i.e. training is stopped when the performance on validation data does not improve, to avoid overfitting. To evaluate the results on the validation set, for regression we used the mean absolute error (MAE) and for classification we used the F1-score. We also used plateau detection to incrementally reduce the learning rate to enhance the optimisation. The initial learning rate was 0.001 and RMSprop (rmsprop) optimizer was used to optimize the network.

Moreover, RF and XGBoost were both tuned using the training dataset, for the different classification and regression tasks and for each cohort, by performing 200 iterations of a random search (scikit-learn implementation of RandomSearchCV()). We used the scikit-learn RF implementation and the XGBoost implementation from the conda-forge channel (https://anaconda.org/conda-forge/xgboost). Supplementary Table 1 reports the best hyper-parameters selected for RF and XGBoost and the settings for the random search. Once tuned on the training data, the three models (RF, XGBoost and NN) were applied to test data. Performances on the test set were compared by looking at the lowest MAE for regression tasks (corneometer and age prediction) and the highest F1-score for classification tasks (smoking and menopausal status).

#### Cross validation

To further examine the stability and the generalisability of our models on unseen data we implemented 5-fold cross validation (5-CV). Table 2 and Supplementary Figure 6,8 summarise the predictive performance of 5-CV using the entire set of samples for Canada cohort and UK cohort. For the Canada cohort we performed cross validation by subject and cross validation using only 1 sample for each subject. Cross validation by subject consists of training each model with the samples from 62 subjects and subsequently making predictions with the trained model on the unseen samples of the 63^rd^ subject. This process is repeated 63 times. See Supplementary Figure 14(a) for performance results. To perform cross validation using 1 sample per subject, we removed the only subject in peri-menopausal status and we then picked the first sample taken, i.e., at the earliest date, for each subject. As such we selected a total of 62 samples. Using the latter subset, we trained and tuned RF and XGBoost to predict the four phenotypes and we performed 5-CV. We did not use the deep NN in this case as a training dataset size of ∼50 is not big enough to report a fair comparison in predictive performance between a deep NN and the other two tree-based ML models. Performance results for RF and XGBoost are shown in Supplementary Figure 14(b).

## Glossary of Terms

AI: artificial intelligence
CV: cross validation
EAI: explainable artificial intelligence
ML: machine learning
MLP: multilayer perceptron
NN: neural network
OTUs: Operational Taxonomy Unit
RF: random forest
SHAP: SHapley Additive exPlanations

## Declarations

### Ethical approval and consent to participate

Written informed consent was obtained from all enrolled individuals. The study protocol was reviewed and approved by Chesapeake IRB, an independent ethics committee. The methods and protocol were carried out in accordance with the approved ICH/GCP guidelines.

### Consent for publication

All authors have consented to this publication.

### Data availability

The data will be made publicly available upon publication.

### Authors’ Contribution

AC conceived the project. NH contributed to the project design, data analysis and biological insights. AC and SM made technical contributions to the machine learning, data analysis and biological insights. LJG made contributions to data analysis and biological insights. EPK made technical contributions to the machine learning. MH, BM, SH, AM, JT and JK provided the data and contributed to the data analysis and biological understanding. CS contributed to the machine learning. MW, WR and LP made contributions to the biological understanding and insights. All authors contributed to the preparation and writing of the manuscript.

#### Acknowledgments

We thank Dr Fausto Martelli, Dr Jason Crain, Dr Nicolas Galichet and Dr Martyn Spink for their support, the useful discussions and their valuable advises.

## Authors’ information

Not applicable

